# Natural compounds magnolol and dicoumarol enhance adipogenesis for future food applications

**DOI:** 10.64898/2026.07.06.736869

**Authors:** Qingwen Xie, N. Stephanie Kawecki, Kathleen K. Chen, Ceci A. Cohen, Emily Cheng, Montgomery Blencowe, Xia Yang, Robert Damoiseaux, Amy C. Rowat

**Affiliations:** Department of Bioengineering, University of California, Los Angeles, Los Angeles, CA 90095, USA; Department of Integrative Biology and Physiology, University of California, Los Angeles, Los Angeles, CA 90095, USA; Department of Chemical and Biomolecular Engineering, University of California, Los Angeles, Los Angeles, CA 90095, USA; Eli and Edyth Broad Stem Cell Center, University of California, Los Angeles, Los Angeles, CA 90095, USA; Department of Molecular and Medical Pharmacology, David Geffen School of Medicine, University of California, Los Angeles, Los Angeles, CA 90095, USA; Institute for Quantitative and Computational Biosciences, University of California, Los Angeles, Los Angeles, CA 90095, USA; California NanoSystems Institute, University of California, Los Angeles, Los Angeles, CA 90095, USA; Jonsson Comprehensive Cancer Center, University of California, Los Angeles, Los Angeles, CA 90095, USA; Molecular Biology Institute, University of California, Los Angeles, Los Angeles, CA 90095, USA

## Abstract

Edible adipose tissue can enhance the sensory and nutritional qualities of cultivated and plant-based meats, yet efficient adipogenic differentiation remains a major bottleneck and synthetic PPARγ agonists are not approved for use in food production. Here, we report a natural compound screen in 3T3-L1 adipocytes that identifies magnolol and dicoumarol as enhancers of adipogenesis; this combination also robustly promotes lipid accumulation in primary porcine dedifferentiated fat cells and ovine preadipocytes. Transcriptomic analyses show that magnolol and dicoumarol induce adipogenesis in murine and porcine cell systems through canonical adipogenic pathways with a narrower transcriptional footprint than the potent PPARγ agonist rosiglitazone. These findings support the potential of naturally occurring compounds magnolol and dicoumarol as enhancers of adipogenesis for both mechanistic studies and food-relevant applications. More broadly, our findings establish a generalizable screening framework and identify small-molecule combinations that accelerate adipose tissue engineering across murine, porcine, and ovine culture systems.

## INTRODUCTION

Future foods – including plant-based, fermentation-derived, and cell-cultured products – have potential as delicious and nutritious complements to animal products. Comparative assessments suggest these approaches can reduce land use and greenhouse gas emissions relative to conventional animal agriculture, contributing to more sustainable food systems^1^. While significant progress has been made in developing protein-rich components of future foods, including engineering skeletal muscle for cell-cultured (or cultivated) meats^2^, fat is central to both food sensory attributes, such as flavor, texture, and mouthfeel, and nutrient composition^3^. Engineered adipose tissue thus has potential to enhance the sensory and nutritional properties of both plant-based and cultivated meats. Beyond food applications, adipocytes are central to metabolic health, making efficient *in vitro* adipogenesis valuable for both fundamental studies of fat cell biology and translational research^4^. Despite the broad importance of adipose tissue, efficiently generating mature, lipid-laden adipocytes *in vitro* remains a challenge.

A major bottleneck in engineering adipose tissue is achieving efficient, robust adipogenic differentiation of preadipocytes. Current protocols rely on synthetic inducers, including 3-isobutyl-1-methylxanthine (IBMX), dexamethasone, insulin, and rosiglitazone^5–11^. These inducers converge on activation of peroxisome proliferator-activated receptor gamma (PPARγ), which is a master regulator of adipogenesis^12^, with the pharmaceutical rosiglitazone acting as a direct PPARγ agonist; however, synthetic inducers carry regulatory and safety limitations that constrain their use in food applications^13,14^. Although efforts to reduce media complexity have established serum-free adipogenic media for multiple species including cow, sheep, pig, and mouse preadipocytes, these approaches still require rosiglitazone, insulin, and species-specific optimization, which limits downstream applicability for food applications^15^.

Naturally occurring compounds represent an underexplored source of adipogenic enhancers that could serve as alternatives to synthetic PPARγ agonists. Dietary lipids such as oleic acid can activate PPARγ and promote lipid accumulation in murine and bovine adipocytes^16–18^, and plant-derived oils such as soybean oil can also induce adipogenesis in mammalian cells^6,19^. More broadly, natural bioactive compounds – including polyphenols, saponins, terpenoids, and fermented plant extracts – show promising effects on activating key adipogenic transcriptional regulators such as PPARγ, C/EBPα, and SREBP1c^20,21^. While natural compounds have potential as adipogenic enhancers for food applications, previous studies have largely focused on testing natural compounds in murine systems, and their efficacy in livestock cells remains largely untested. Murine 3T3-L1 preadipocytes provide a well-established model for studying adipogenesis^4,22^, and while previous studies have shown translatability of findings to edible species based on phenotypic observations, mechanistic characterizations across cell systems of different species have been lacking; yet comparative transcriptomic and proteomic studies across species are necessary to develop predictive understanding of adipogenic responses in murine models that can translate to other food-relevant species. Moreover, most studies test individual compounds in isolation, despite evidence that adipogenic induction protocols typically require multi-component cocktails to achieve robust differentiation^17,19,23^. Currently, there is no systematic or generalizable experimental framework for identifying combinations of natural compounds that can reliably accelerate adipogenesis in both model systems and food-relevant species.

Here, we developed a systematic screening approach to identify naturally derived small molecules that enhance adipogenesis. We reasoned that natural bioactive compounds could offer practical advantages over synthetic PPARγ agonists for food-relevant applications, and screened libraries of ∼5000 naturally derived compounds in murine 3T3-L1 preadipocytes to identify novel adipogenic enhancers. The screen identified magnolol and dicoumarol as enhancers of adipogenesis, and we demonstrate that the combination of these natural compounds accelerates adipogenesis in primary dedifferentiated porcine fat (DFAT) cells and ovine preadipocytes. Integrated multi-omics analysis reveals that magnolol and dicoumarol activate core adipogenic pathways while inducing a narrower transcriptional footprint than rosiglitazone. Together, these findings establish both a screening strategy and a mechanistic framework for developing food-compatible adipogenic media and scalable biomanufacturing of adipose tissue for sustainable food applications.

## RESULTS

### Screen of natural compounds identifies dicoumarol and magnolol as adipogenic enhancers

To build a systematic framework for identifying adipogenic enhancers that can translate from murine models to food-relevant species, we developed the screen to detect increases in lipid accumulation in BODIPY-stained 3T3-L1 murine preadipocytes (**Figure 1A**). As a positive control to validate assay sensitivity and dynamic range, we used the PPARγ agonist, rosiglitazone (**Figure 1B**). The assay was developed to have a Z’ ≥ 0.5, the standard threshold for high-throughput screening assay validation^24^.

**Figure 1.**
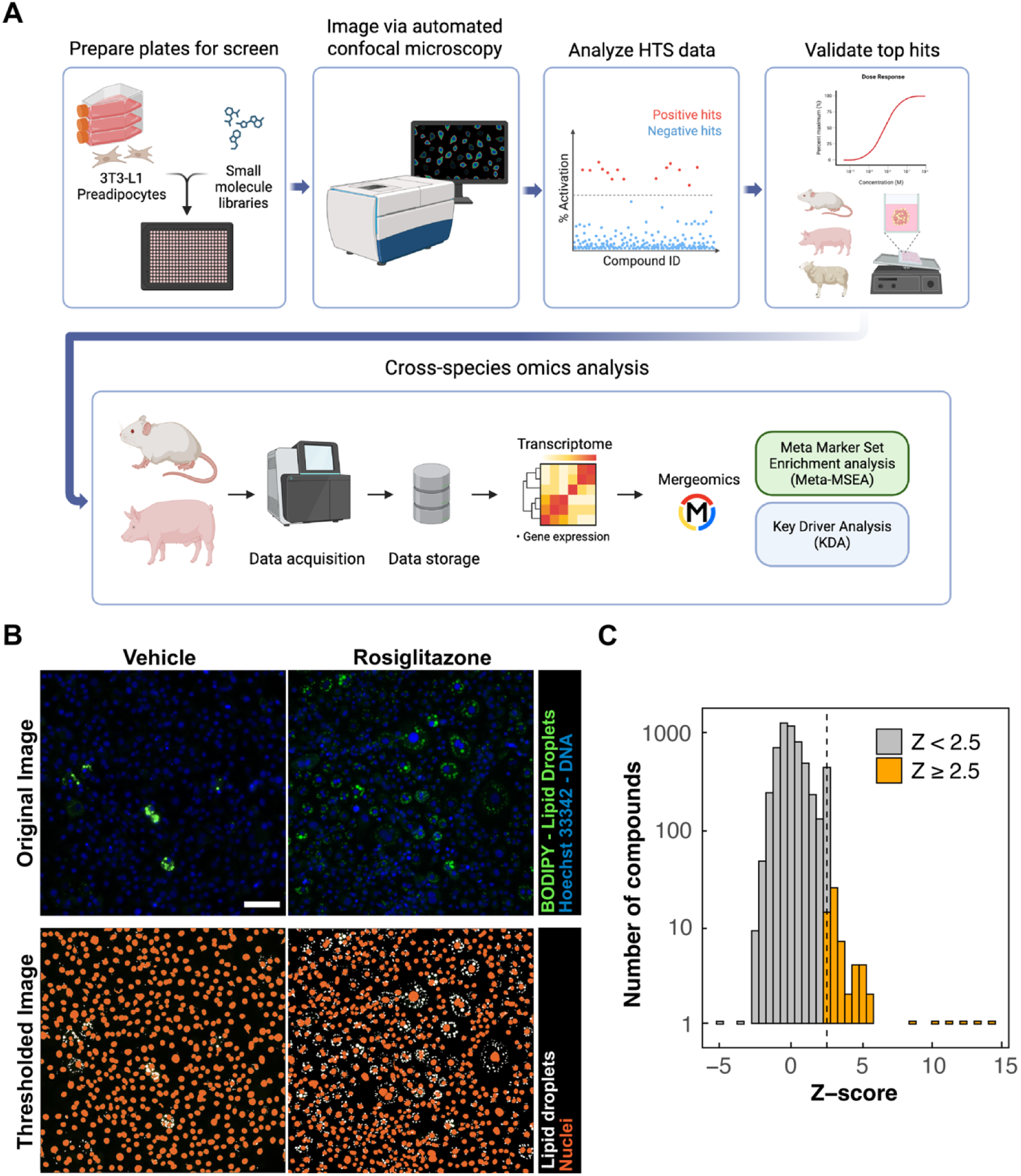
Identifying adipogenic enhancers using a high-throughput screen (HTS). (A) Schematic diagram of the HTS to identify enhancers of adipogenesis. (B) Examples of the merged fluorescent images captured by automated confocal microscopy. Lipids are stained by BODIPY 493/503, and DNA by Hoechst 33342. Scale, 100 µm. (C) Distribution of Z-scores (Z) across the ∼ 5,000 compounds from Spectrum Collection, Natural Organic Compound Library, and Natural Product Library assayed in the screen (n = 1). We defined hits as compounds that results in increased intracellular lipid area of more than 2.5 standard deviations from the mean. Gray bars indicate compounds with Z < 2.5; orange bars indicate compounds with Z ≥ 2.5. Dashed vertical line delineates Z = 2.5.

To identify novel enhancers of adipogenesis, we treated 3T3-L1 murine preadipocytes with ∼5,000 natural compounds, imaged the plates after 4 days of differentiation, and calculated a Z-score for each compound based on the projected area of intracellular lipids per cell (**Figure 1C**). To identify compounds with adipogenic activity that was significantly higher than the population mean, we selected compounds with Z-score > 2.5; this threshold is commonly used in high-throughput screening for hit selection^25^. We x compounds as hits with Z-scores > 2.5 (**Figure 1C**).

Dicoumarol and magnolol were prioritized for follow-up studies based on their strong adipogenic activity, favorable viability profiles, and status as naturally derived bioactive compounds^26,27^ (**Figure 2A**). Dicoumarol derives from sweet clover, is a vitamin K antagonist, and has been used as an anticoagulant^26^. Magnolol is isolated from *Magnolia* species, has been used in traditional herbal medicine and as a dietary supplement^27,28^, and explored as a dietary feed supplement for livestock^29–32^. We first investigated the dose-dependent effects of dicoumarol and magnolol on 3T3-L1 adipocytes. Treatment with increasing concentrations of either dicoumarol or magnolol resulted in increased lipid accumulation (**Figures 2B–D**). While both compounds have been reported to induce apoptosis in cancer cells^26,33^, we found no significant reduction in the viability of 3T3-L1 adipocytes at concentrations up to 10 µM (**Figure 2E**), making them suitable candidates as adipogenic enhancers.

**Figure 2.**
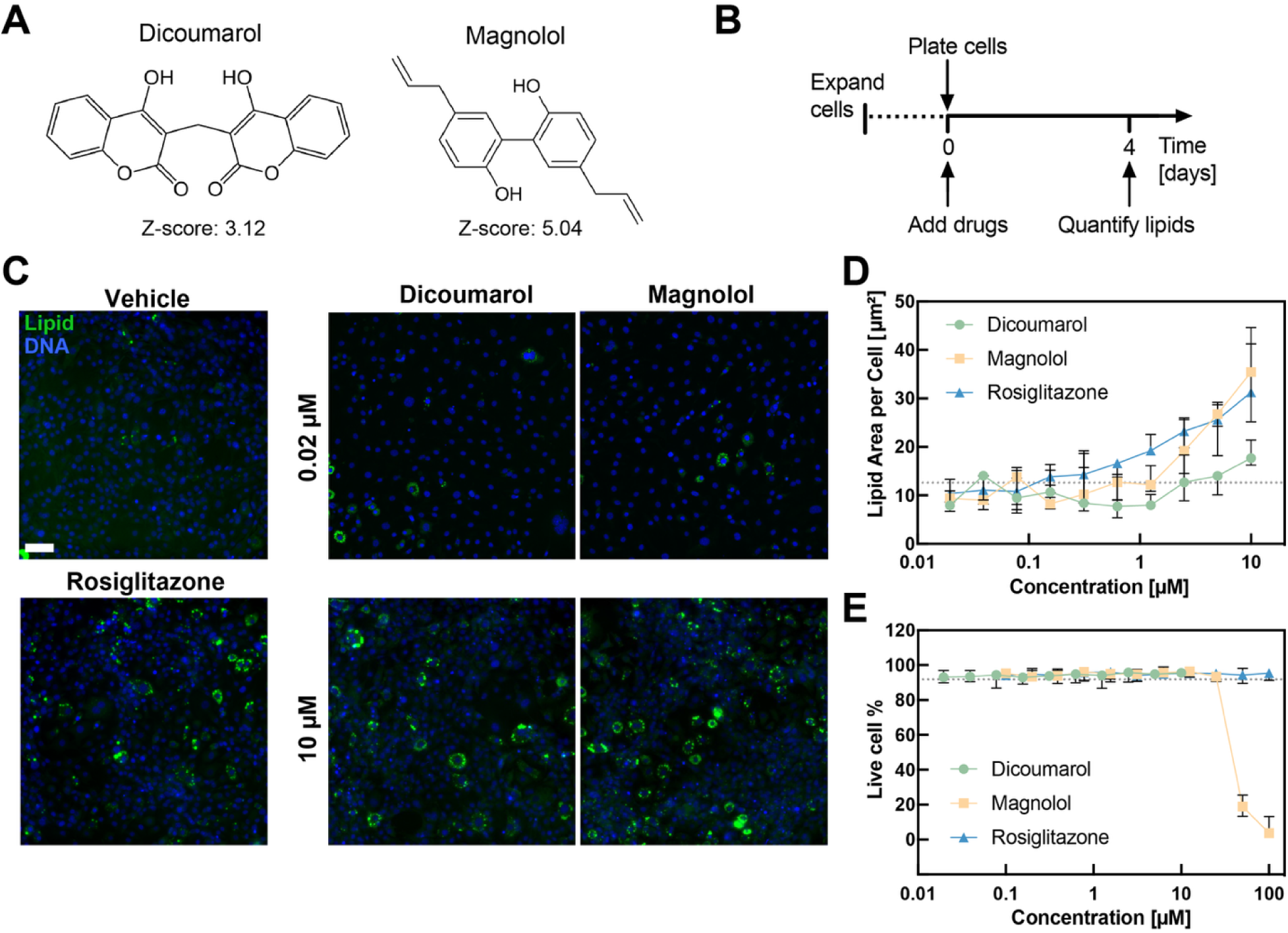
Top hits dicoumarol and magnolol increase lipid accumulation in murine 3T3-L1 adipocytes. (A) Chemical structures of the two top hits, dicoumarol and magnolol. (B) Experimental timeline for assessing adipogenesis in murine 3T3-L1 adipocytes using quantitative image analysis. (C) Representative confocal images of 3T3-L1 adipocytes after 4 days of adipogenesis with treatments of dicoumarol, magnolol, rosiglitazone, and vehicle (DMSO) control. Intracellular lipids are visualized using BODIPY (green); DNA is stained using Hoechst 333342 (blue). Scale, 100 μm. (D) Quantification of lipid area in 3T3-L1 adipocytes treated with dicoumarol, magnolol, rosiglitazone, and vehicle (DMSO) control (n = 3 independent experiments, total ∼10,800 individual cells per experiment). Quantitative image analysis is conducted on confocal images acquired at 4 days after treatment with compound/vehicle; findings are normalized to the number of cells, as determined by the nuclear count. Dotted line indicates the vehicle-treated baseline. (E) Percentage of live cells after 4-day treatment with dicoumarol, magnolol, and rosiglitazone determined by Calcein-AM/Propidium Iodide Live/Dead staining (n = 3 independent experiments, total ∼6000 individual cells per experiment). Dotted line indicates the vehicle-treated baseline.

To test whether the effects of dicoumarol and magnolol were robust across adipogenic induction protocols, we evaluated them using a reduced MDI medium, which is a simplified version of the widely used MDI adipogenic induction medium (M - IBMX, D - dexamethasone, and I - insulin). Consistent with our screening results, 3T3-L1 cells differentiated with reduced MDI medium showed a significant increase in lipid accumulation when treated with magnolol (**Figures 3A–3C**). Dicoumarol also induced increased lipid accumulation compared to vehicle control, although the effect was more subtle than magnolol. These findings demonstrate that the effects of dicoumarol and magnolol replicate across distinct differentiation protocols.

**Figure 3.**
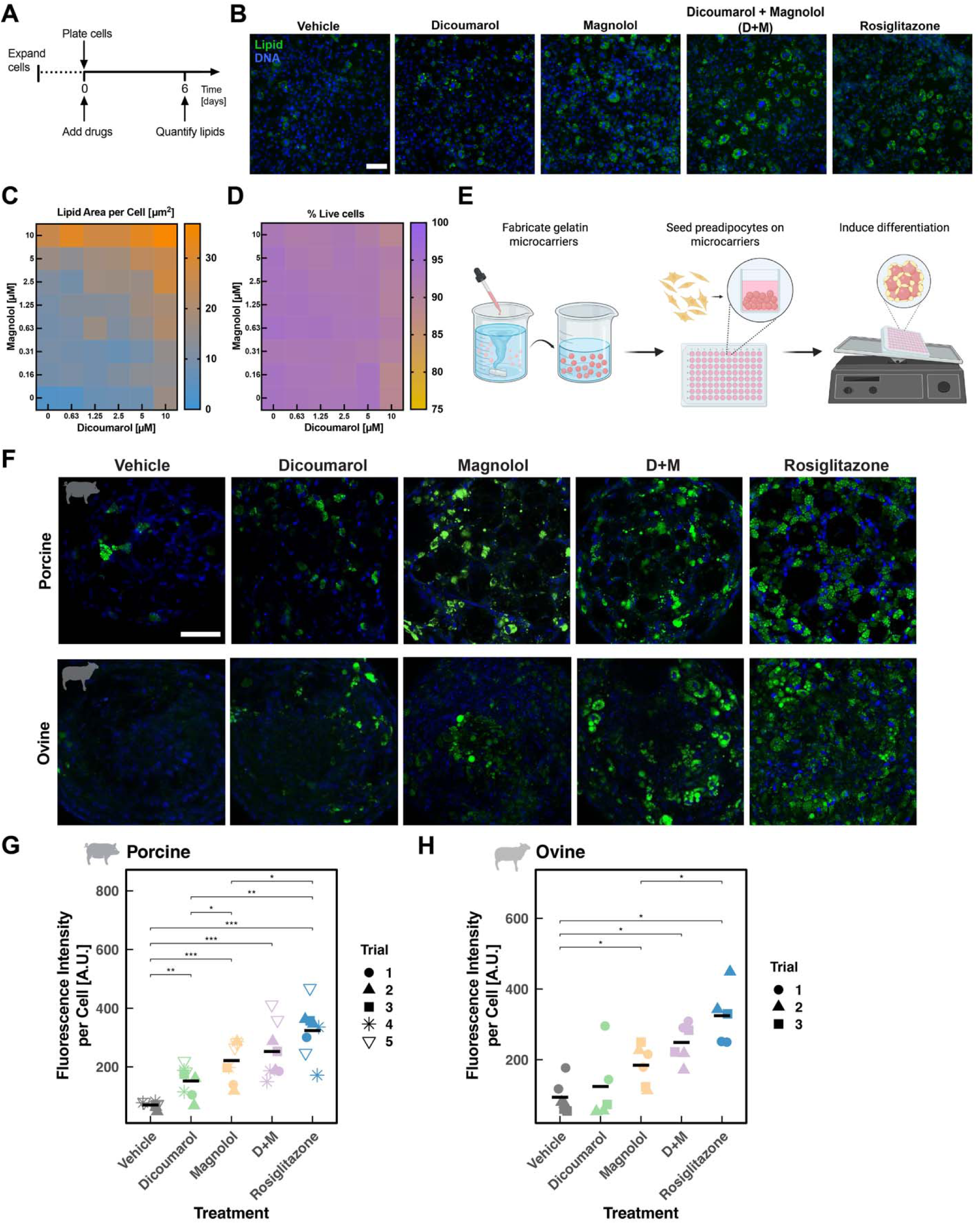
Combined effects of dicoumarol and magnolol. (A) Experimental timeline for assessing adipogenesis in 3T3-L1 adipocytes using quantitative image analysis. (B) Lipid-stained (BODIPY, green) and DNA-stained (Hoechst 333342, blue) images of 3T3-L1 adipocytes after 6 days of adipogenesis. Scale, 100 μm. (C) Heatmap representation of the synergistic interactions between dicoumarol and magnolol in 3T3-L1 adipocytes. Results show the median values from n = 4 independent experiments (total ∼10,800 individual cells per condition). (D) Heatmap showing cell viability across dicoumarol and magnolol treatments assessed by Calcein-AM/propidium iodide staining. Results show the mean values from n = 3 independent experiments (total ∼6,000 individual cells per condition). Color scale indicates the percentage of live cells to total cells from low (blue) to high (orange). (E) Schematic of the edible microcarrier-based culture system. (F) Lipid and DNA staining of porcine and ovine preadipocytes differentiated on edible microcarriers for 7 days. Cells are stained for lipids using BODIPY (green) and for DNA using Hoechst 33342 (blue). Scale, 100 μm. (G-H) Quantification of fluorescence intensity per cell in (G) porcine (n = 5 independent experiments, total ∼400 individual cells per condition) and (H) ovine (n = 3 independent experiments, total ∼200 individual cells per condition) preadipocytes based on BODIPY and Hoechst 33342 staining. Different symbols represent independent experiments. Statistical comparisons are shown relative to vehicle (*p<0.05, **p<0.01, ***p<0.001) using Mann-Whitney U test.

We next tested if treatment with both dicoumarol and magnolol (D+M) had any combined effects on adipogenesis. Interestingly, we found that D+M combinations across a range of concentrations up to 10 µM strongly enhanced lipid accumulation compared to either compound alone, with the most significant effects at ≥ 5 µM magnolol and ≥ 5 µM dicoumarol (**Figures 3A–3C**). Since increasing concentrations of both dicoumarol and magnolol could have cytotoxic effects, we assessed the viability of cells treated with dicoumarol and magnolol using Calcein-AM/propidium iodide staining; this revealed no significant changes in the viability of 3T3-L1 cells across the full range of compound combinations and concentrations compared to vehicle-treated cells (**Figure 3D**). These findings demonstrate that the combined D+M treatment can enhance adipogenesis compared to either compound alone.

### Combined treatment of dicoumarol and magnolol enhances adipogenesis in porcine and ovine adipocytes with reduced dependence on synthetic adipogenic inducers

To determine if our findings are translatable to edible cell types in a scalable culture system, we next tested the effects of D+M on porcine DFAT cells and ovine pre-adipocytes cultured on edible microcarriers in suspension culture. In these culture conditions, we previously showed that 3T3-L1 adipocytes on microcarriers adhered to each other together to form multicellular aggregates or microtissues^34,35^. When cultured with edible microcarriers, we found that porcine DFAT cells in microtissues showed a significant increase in lipid accumulation upon treatment with magnolol (**Figures 3E and 3F**), and a modest increase with dicoumarol; these findings are consistent with observations of increased lipid accumulation in 3T3-L1 treated with dicoumarol and magnolol in 2D cultures (**Figure 2C**). The D+M combination treatment further enhanced lipid accumulation in porcine DFAT cells on edible microcarriers^35^ (**Figures 3E and 3F**), validating the combined effect of magnolol and dicoumarol in an edible cell type.

Ovine pre-adipocytes differentiated on edible microcarriers showed a similar increase in lipid accumulation with D+M treatment compared to vehicle control (**Figures 3F and 3H**). Importantly, D+M treatment showed a combined effect even without IBMX, whereas vehicle-treated cells failed to differentiate under the same conditions; these findings suggest that dicoumarol and magnolol can reduce the dependence of this commonly used synthetic adipogenic inducer, which has oral toxicity and no regulatory approval for human consumption^36^. These findings support the potential of dicoumarol and magnolol to increase the efficacy of edible adipose tissue growth in a scalable, suspension-based format with food-relevant cell types.

### Dicoumarol and magnolol activate core adipogenic pathways with a narrower transcriptional footprint than rosiglitazone

To investigate the molecular mechanisms underlying the enhanced adipogenesis of dicoumarol and magnolol, we used omics approaches to measure changes in transcript levels in 3T3-L1 adipocytes treated with the compounds. We first examined transcriptional changes after 5 days of differentiation using bulk RNA-sequencing (RNA-seq) to identify differentially expressed genes (DEGs) between each treatment and vehicle (DMSO) control at a false discovery rate FDR < 0.05. Volcano plots of DEGs for each treatment – dicoumarol, magnolol, combination D+M treatment, and the positive control, rosiglitazone – showed that dicoumarol alone induced minimal transcriptional changes (1 upregulated DEG), while magnolol had 245 upregulated and 59 downregulated DEGs, and the combination of D+M showed 730 upregulated and 344 downregulated DEGs (**Figures 4A–4D**). By contrast, rosiglitazone had approximately 4-10 times as many DEGs with 3,310 upregulated and 3,180 downregulated genes. Although rosiglitazone was used at a lower concentration (1 µM vs. 10 µM for dicoumarol and magnolol), it is a potent and selective PPARγ agonist^38^, which likely accounts for its larger transcriptional footprint.

**Figure 4.**
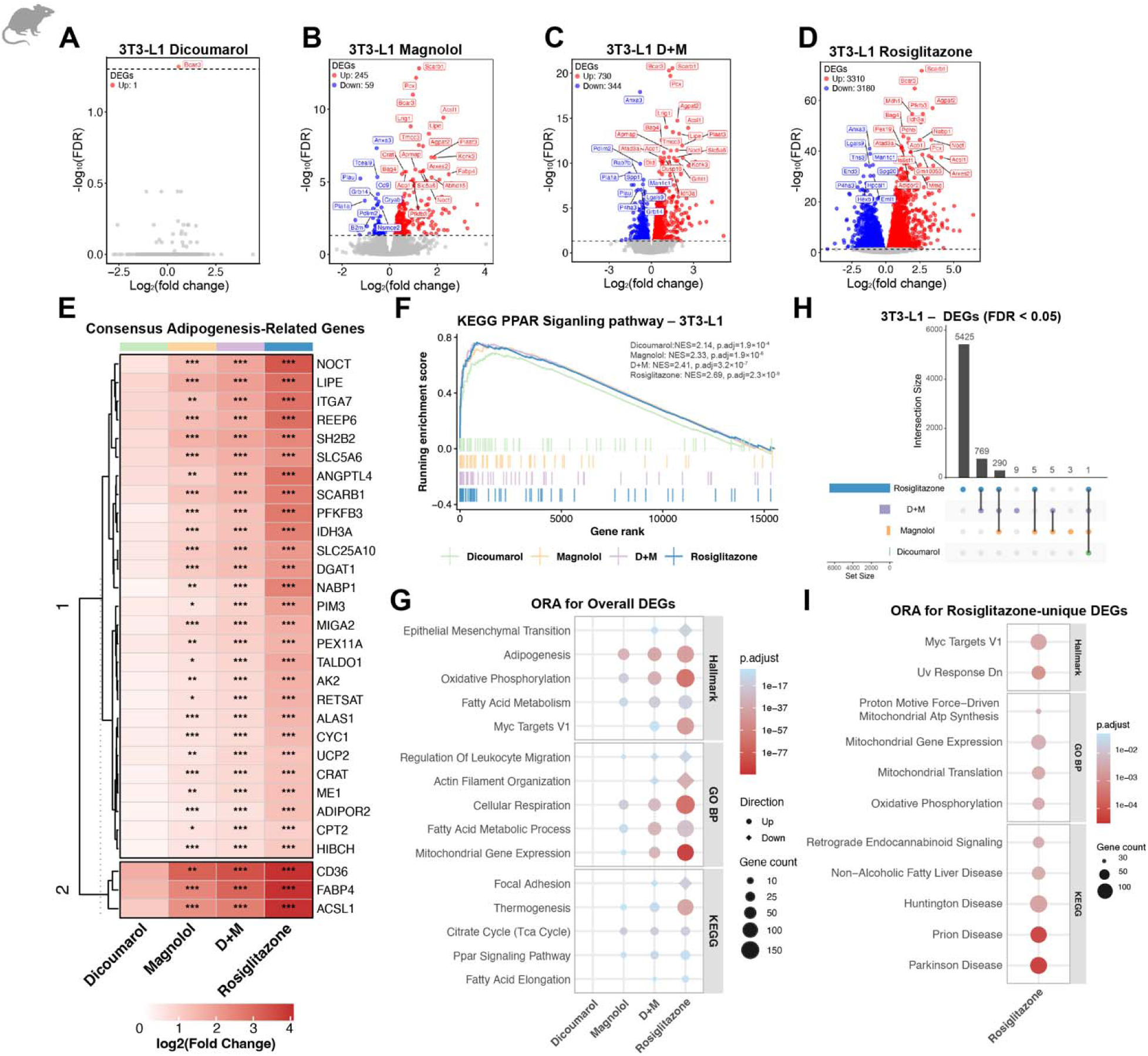
Transcriptomic changes induced by dicoumarol and magnolol in 3T3-L1 adipocytes. (A–D) Volcano plots to visualize differentially expressed genes (DEGs; FDR < 0.05) in 3T3-L1 adipocytes treated with: (A) dicoumarol, (B) magnolol, (C) dicoumarol and magnolol (D+M), or (D) rosiglitazone compared to vehicle-treated controls. Red dots indicate upregulated DEGs; blue dots downregulated DEGs. Total counts of upregulated (up) and downregulated (down) DEGs are indicated in each panel. (E) Heatmap of the top adipogenesis-associated DEGs across all four treatments, showing log_2_(fold change) relative to vehicle. Shown in the heatmap are 30 genes out of 349 adipogenesis-related murine genes queried. Genes are grouped into two clusters based on unsupervised hierarchical clustering: Cluster 1 comprises genes with moderate upregulation across treatments; Cluster 2 comprises canonical adipogenic markers showing the strongest upregulation. Asterisks indicate statistical significance: *p_adj_ < 0.05, ** p_adj_ < 0.01, ***p_adj_ < 0.001. (F) Gene set enrichment analysis (GSEA) enrichment plot for the KEGG PPAR signaling pathway gene set in 3T3-L1 adipocytes. Running enrichment scores, normalized enrichment scores (NES), and p_adj_ values are shown for dicoumarol (green), magnolol (orange), D+M (purple), and rosiglitazone (blue). (G) Overrepresentation analysis (ORA) dot plot showing selected enriched pathways across Hallmark, GO Biological Process (GO BP), and KEGG categories for all four treatments. Pathways shown represent non-redundant terms from each database, selected based on statistical significance (p_adj_ < 0.05) and largest gene counts, with redundant parent-child GO terms removed using semantic similarity filtering. Dot size indicates gene count; color indicates p_adj_; and shape indicates upregulated (circle) or downregulated (diamond) gene sets. (H) UpSet plot showing the shared and unique DEGs across dicoumarol, magnolol, D+M, and rosiglitazone treatments. (I) ORA dot plot of top enriched pathways in the 5,425 rosiglitazone-unique DEGs across Hallmark, GO BP, and KEGG categories. Top pathways were selected based on statistical significance (p_adj_ < 0.05) and prioritized for the highest enrichment scores and largest gene counts, with redundant parent-child GO terms removed using semantic similarity filtering. No significant pathway enrichment was detected for dicoumarol, magnolol, or dicoumarol and magnolol (D+M).

To map how these compounds impact adipogenesis at the transcriptional level, we curated a panel of consensus adipogenesis-associated genes from three complementary gene set databases: Hallmark Adipogenesis, Gene Ontology: Biological Process (GO: BP) Fat Cell Differentiation, and Reactome PPAR Signaling (349 mouse genes in total). We selected the top 30 genes that had p_adj_ < 0.5 across the maximum number of treatments and an absolute log_2_FC > 0.5 in at least one condition. A heatmap of these DEGs showed consistent up-regulation across two clusters (**Figure 4E**). Cluster 2 showed stronger upregulation of canonical adipogenic markers including *Fabp4, Cd36, Acsl1,* with fold changes following the trend of the phenotypic lipid accumulation results (**Figure 2 and Figure 3**): dicoumarol < magnolol < D+M < rosiglitazone. Cluster 1 genes such as *Adipor2, Cpt2, Me1, Ucp2* showed more moderate expression changes following the same trend (**Figure 4E**). Because PPARγ is a key regulator of adipogenesis, we evaluated PPAR signaling using the threshold-free GSEA enrichment analysis; this confirmed significant enrichment of the Kyoto Encyclopedia of Genes and Genomes (KEGG) PPAR signaling pathway gene set for all four treatments (**Figure 4F**; all p_adj_ <0.05). Notably, despite dicoumarol producing only one significant DEG, GSEA revealed positive enrichment of adipogenesis-related genes; this suggests that dicoumarol induces a shift in cells towards an adipogenic state through modest, coordinated transcriptional changes rather than driving large expression changes at the individual gene level.

To assess transcriptional changes more broadly, we used overrepresentation analysis (ORA) of DEGs against Hallmark, GO:BP, and KEGG gene sets. For visualization, we selected representative, non-redundant terms from each database, prioritizing terms with the strongest statistical significance and largest gene counts while removing highly redundant parent-child GO terms and disease-associated KEGG pathways enriched due to gene set annotation overlap. While dicoumarol showed no significant enrichment as it only had one significant DEG, we found similar enrichment trends for magnolol, D+M, and rosiglitazone (**Figure 4G**). Hallmark pathway analysis revealed significant enrichment of adipogenesis, oxidative phosphorylation, and fatty acid metabolism, while KEGG pathway analysis identified enrichment of thermogenesis, the tricarboxylic acid (TCA) cycle, and PPAR signaling. GO:BP analysis showed enrichment of processes central to adipocyte differentiation and lipid accumulation including cellular respiration, fatty acid metabolic processes, and mitochondrial gene expression. Downregulated pathways included epithelial-mesenchymal transition (EMT), reflecting loss of preadipocyte mesenchymal identity during adipogenic commitment, and regulation of leukocyte migration – a gene set that includes inflammatory chemokines and cytokines, whose downregulation is consistent with the reduced inflammatory signaling for the promotion of adipogenesis^37^. With the exception of dicoumarol, all treatments followed a consistent trend across all pathway categories, but the magnitude of enrichment varied: magnolol alone showed moderate enrichment, D+M showed stronger enrichment, and rosiglitazone showed the broadest and most significant enrichment; this pattern mirrors the phenotypic lipid accumulation data showing increased lipid accumulation (**Figure 2 and Figure 3**) and supports that D+M activates similar adipogenic programs as rosiglitazone. The similar trend across D+M and rosiglitazone treatments was also reflected in the downregulated pathways, actin filament organization and focal adhesion. The suppression of actin and focal adhesion-associated genes may reflect the cytoskeletal remodeling and loss of cell-matrix interactions that accompany the transition from an elongated preadipocyte to a rounded adipocyte morphology during adipogenic commitment^38–40^.

To further evaluate similarities and differences in transcriptional signatures across treatments, we examined the overlapping and unique DEGs (**Figure 4H**). Magnolol, D+M, and rosiglitazone shared 290 DEGs including canonical adipogenic markers such as *Fabp4, Adipoq, Lipe, Plin2*, and *Acsl1*; these findings are consistent with the observation that all compound treatments promote adipogenesis (**Figure 4E**). Analysis of the unique DEGs showed that dicoumarol and magnolol showed very few unique DEGs: dicoumarol had none, magnolol had 3, and D+M had 9. Accordingly, we did not identify any significant pathway enrichment for these genes through ORA, indicating that the unique transcriptional programs induced do not converge on specific biological processes. By contrast, rosiglitazone uniquely regulated 5,425 genes that were not shared with the natural compound treatments. To characterize the biological significance of rosiglitazone’s unique transcriptional programs, we analyzed pathways enriched in these DEGs (**Figure 4I**); this revealed significant enrichment in mitochondrial gene expression, mitochondrial translation, and oxidative phosphorylation (OXPHOS), as well as MYC targets and UV responses. While several KEGG pathways – including retrograde endocannabinoid signaling, non-alcoholic fatty liver disease, Huntington’s disease, Prion disease, and Parkinson’s disease – were significantly enriched in rosiglitazone-unique genes, these pathways have significant overlap with mitochondrial OXPHOS gene sets and the enrichment likely reflects mitochondrial activity rather than disease-specific mechanisms. Beyond mitochondrial processes, enrichment of MYC targets reflect that rosiglitazone also activates proliferative and biosynthetic programs; this is consistent with the role of PPAR agonists in regulating adipogenesis, including a proliferative phase that precedes terminal adipocyte differentiation, as well as ribosome biogenesis, which supports the increased biosynthetic demands of lipid accumulation^41^. Taken together, these findings demonstrate that while D+M and rosiglitazone similarly activate core adipogenic programs, rosiglitazone drives a substantially broader adipogenic response that includes distinct mitochondrial, metabolic, and proliferative programs.

While murine 3T3-L1 cells provide a well-established model for adipogenesis, we next asked how these findings would translate to a cell type with stronger relevance for food applications by conducting bulk RNA-seq on porcine DFAT cells. Volcano plots revealed that dicoumarol induced the fewest transcriptional changes (179 upregulated, 241 downregulated DEGs), while magnolol produced a more substantial response (570 upregulated, 305 downregulated) (**Figures 5A–5D**). The combination of D+M further amplified the transcriptional response (869 upregulated, 871 downregulated), which was comparable in magnitude to rosiglitazone (970 upregulated, 828 downregulated). This pattern is broadly consistent with our mouse findings, although D+M and rosiglitazone showed similar magnitudes of DEGs in porcine cells, whereas the transcriptional changes induced by rosiglitazone were substantially larger in mouse.

**Figure 5.**
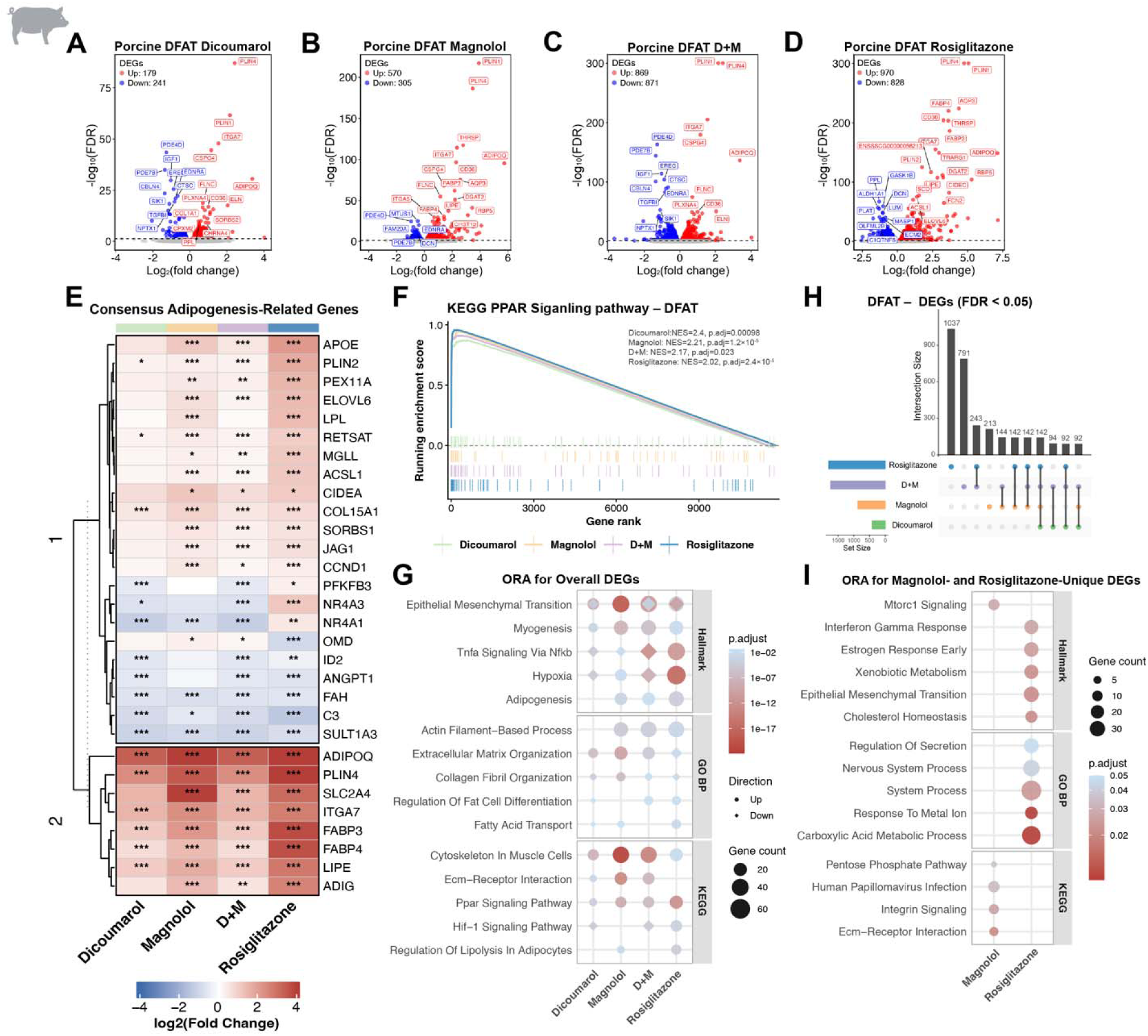
Dicoumarol and magnolol show adipogenic effects in DFAT at the transcriptional level. (A–D) Volcano plots to visualize differentially expressed genes (DEGs; FDR < 0.05) in porcine DFAT adipocytes treated with: (A) dicoumarol, (B) magnolol, (C) dicoumarol and magnolol (D+M), or (D) rosiglitazone compared to vehicle-treated controls. Red dots indicate upregulated DEGs; blue dots downregulated DEGs. Total counts of upregulated (up) and downregulated (down) DEGs are indicated in each panel. (E) Heatmap of the top adipogenesis-associated DEGs across all four treatments, showing log_2_(fold change) relative to vehicle. Shown in the heatmap are 30 genes out of 341 adipogenesis-related porcine genes queried. Genes are grouped into two clusters based on unsupervised hierarchical clustering: Cluster 1 comprises genes with moderate up- or down- regulation across treatments; Cluster 2 comprises canonical adipogenic markers showing the strongest upregulation. Asterisks indicate statistical significance: *p_adj_ < 0.05, ** p_adj_ < 0.01, ***p_adj_ < 0.001. (F) Gene set enrichment analysis (GSEA) enrichment plot for the KEGG PPAR signaling pathway gene set in DFAT cells. Running enrichment scores, normalized enrichment scores (NES), and p_adj_ values are shown for dicoumarol (green), magnolol (orange), D+M (purple), and rosiglitazone (blue). Running enrichment scores are shown for dicoumarol (green), magnolol (orange), D+M (purple), and rosiglitazone (blue). Normalized enrichment scores (NES) and p_adj_ values for dicoumarol: NES=2.4, p_adj_ = 0.00098; Magnolol: NES=2.21, p_adj_ = 1.2 × 10^-5^; D+M:NES=2.17, p_adj_ = 0.023; Rosiglitazone: NES=2.02, p_adj_ = 2.4 × 10^-5^. (G) Overrepresentation analysis (ORA) dot plot showing selected enriched pathways across Hallmark, GO Biological Process (GO BP), and KEGG categories for all four treatments. Pathways shown represent non-redundant terms from each database, selected based on statistical significance (p_adj_ < 0.05) and largest gene counts, with redundant parent-child GO terms removed using semantic similarity filtering. Dot size indicates gene count; color indicates p_adj_; and shape indicates upregulated (circle) or downregulated (diamond) gene sets. (H) UpSet plot showing the shared and unique DEGs across dicoumarol, magnolol, D+M, and rosiglitazone treatments. (I) ORA dot plot of top enriched pathways in the 213 magnolol-unique DEGs and 1,037 rosiglitazone-unique DEGs across Hallmark, GO BP, and KEGG categories. Top pathways were selected based on statistical significance (p_adj_ < 0.05) and prioritized for the highest enrichment scores and largest gene counts. No significant pathway enrichment was detected for dicoumarol or dicoumarol and magnolol (D+M).

To evaluate the effects of the natural compounds on adipogenesis in porcine cells, we assessed the expression of the same three gene sets as in **Figure 4E**. The top adipogenesis-associated DEGs showed two distinct expression clusters (**Figure 5E**). Cluster 2 comprised canonical adipogenic markers including the adipokine *ADIPOQ*; lipid-handling genes *FABP4, FABP3, PLIN4, ADIG* and *LIPE;* the glucose transporter *SLC2A4;* and integrin *ITGA7*; these genes were consistently upregulated across magnolol, D+M, and rosiglitazone treatments, confirming the activation of adipogenic programs. Dicoumarol showed a similar trend, although the magnitude of gene expression change was smaller, with *SLC2A4* and *ADIG* not reaching significance. Consistent with these findings, GSEA confirmed significant enrichment of the KEGG PPAR signaling pathway across all four treatments (all p_adj_ < 0.05; **Figure 5F**), supporting that PPARγ activation is a shared mechanism across both murine and porcine adipogenic systems.

Genes in cluster 1 showed more heterogeneous expression patterns. For example, *NR4A1*, a negative regulator of adipogenesis that maintains preadipocyte quiescence, was downregulated by the natural compounds. By contrast, *NR4A1* and *NR4A3* were specifically upregulated by rosiglitazone, consistent with their known induction by thiazolidinedione-class PPARγ agonists and role in promoting insulin sensitivity^42^. Several other cluster 1 genes including *FAH, C3, SULT1A3*, and *ANGPT1* showed downregulation across multiple treatments; these genes are included in adipogenic gene sets as co-expressed adipose tissue markers, which represent metabolic enzymes, complement components, and vascular factors, rather than core adipogenic drivers. Their downregulation in DFAT cells likely reflects loss of the dedifferentiated fibroblast-like state during redifferentiation, rather than suppression of adipogenic programs.

To characterize pathway-level changes more broadly, we performed ORA on all DEGs across treatments (**Figure 5G**). All compounds showed enrichment of PPAR signaling, consistent with their role in promoting adipogenesis. Adipogenesis was enriched across magnolol, D+M, and rosiglitazone, while fatty acid transport showed enrichment for dicoumarol, magnolol, and rosiglitazone. Across all treatments, we also observed enrichment for myogenesis, actin filament-based processes, and cytoskeleton in muscle cells, reflecting the structural reorganization accompanying porcine DFAT cell redifferentiation; this pattern was more prominent in the porcine system and observed to a lesser extent in mouse. ECM organization, collagen fibril organization, and ECM-receptor interaction were all positively enriched in natural compounds treatments but negatively enriched or absent in rosiglitazone, suggesting a divergent ECM regulatory program between dicoumarol and magnolol versus rosiglitazone. EMT also showed strong enrichment across treatments with mixed directionality, where structural ECM-producing genes were upregulated while signaling and regulatory factors were downregulated, likely reflecting the concurrent structural remodeling and transcriptional reorganization that accompanies DFAT cells redifferentiation into adipocytes.

Rosiglitazone showed a distinct upregulated program in TNFα/NF-κB and HIF-1 signaling pathways in porcine cells, indicating potential off-target inflammatory activation and hypoxic pathway activation (**Figure 5G**). By contrast, TNFα/ NF-κB signaling was downregulated across natural compound treatments, indicating suppression of inflammation during adipogenic commitment. HIF-1 signaling was additionally downregulated in dicoumarol and D+M but not magnolol, suggesting that dicoumarol may be the primary driver of HIF-1 suppression in the combination treatment in porcine cells.

Examination of the shared and unique DEGs in porcine cells revealed 142 genes that were common to the natural compound and rosiglitazone treatments (**Figure 5H**). These shared DEGS included core adipogenic regulators such as *ADIPOQ, CD36, FABP4, PLIN1*, and *CIDEC*, supporting conserved activation of adipogenic programs across porcine and murine systems. Analysis of unique DEGs showed that dicoumarol had none while magnolol, D+M, and rosiglitazone uniquely regulated 213, 791, 1037 DEGs, respectively. While D+M uniquely regulated 791 genes, these showed no significant pathway enrichment, indicating that D+M’s adipogenic effects are driven primarily through the 142 shared genes, and the unique D+M DEGs do not converge on a coherent biological program.

Pathway analysis of the unique DEGs in the porcine cell system revealed distinct transcriptional programs for magnolol and rosiglitazone beyond the shared core adipogenic DEGs. Magnolol-unique DEGs showed enrichment in mTORC1 signaling, pentose phosphate pathway, integrin signaling, and ECM-receptor interaction (**Figure 5I**); these pathways may support structural reorganizations that accompany adipogenesis during DFAT redifferentiation. By contrast, rosiglitazone-unique genes showed enrichment in metabolic processes including xenobiotic metabolism, cholesterol homeostasis, and carboxylic acid metabolic process consistent with its broader PPARγ-mediated transcriptional activation and its activity as a pharmaceutical compound. The enrichment of estrogen response in rosiglitazone-unique genes may reflect the known interaction between rosiglitazone and estrogen signaling in adipose tissue^43^. Rosiglitazone-unique DEGs also showed enrichment for interferon gamma response, which supports the broader off-target inflammatory transcriptional activation – consistent with the upregulation of TNFα/NF-κB signaling shown in **Figure 5G**. Interestingly, unique DEGs induced by rosiglitazone show enrichment for EMT; since EMT is also enriched among shared DEGs, these findings substantiate that rosiglitazone engages EMT-related programs more broadly than the natural compounds. These findings suggest that beyond the shared adipogenic programs, rosiglitazone induced more extensive gene expression changes than the natural compounds.

Comparison across murine 3T3-L1 and porcine DFAT cell systems revealed conserved PPAR signaling enrichment and core adipogenic gene induction in response to both magnolol, D+M, and rosiglitazone supporting a shared mechanism of action. Despite inducing fewer total DEGs, all natural compound treatments induced prominent ECM and cytoskeletal gene signatures in porcine cells, which likely reflect the distinct cellular context of porcine DFAT cells undergoing dedifferentiation-redifferentiation, compared to the committed preadipocyte identity of the immortalized murine 3T3-L1 cells. Across both systems, rosiglitazone consistently drove the broadest and strongest transcriptional response in the majority of pathways, whereas magnolol and D+M exhibited more targeted transcriptional profiles centered on core adipogenic programs.

### Key driver analysis identifies regulatory networks underlying adipogenic responses across murine and porcine systems

To identify regulatory networks underlying the transcriptomic changes observed with compound treatments, we used Mergeomics^44^ to infer upstream regulators – or key driver genes – and their associated networks. This analysis revealed both shared and treatment-specific predicted regulatory networks across murine and porcine cell systems.

Consistent with our DEG analysis, the number and connectivity of predicted key drivers reflected the magnitude of transcriptional response for each treatment. In mouse 3T3-L1 adipocytes, predicted key drivers organized into functional clusters including adipogenesis and lipid metabolism, oxidative phosphorylation, TCA cycle, cytoskeleton and cell mechanics, ECM, immune signaling, cell cycle, and translation (**Figure 6A and Figure S1**). The highest-centrality nodes formed a metabolic core enriched for PPARγ target genes, lipid handling, and mitochondrial oxidative metabolism, which was shared across multiple treatments and consistent with the convergent activation of adipogenic metabolic programs. This dominance of metabolic hubs in murine networks reflects the committed preadipocyte identity of 3T3-L1 cells. Key drivers of this metabolic core included canonical adipogenic regulators, such as FASN and SCD (**Figure S1**). Within the core themes, we also observed that some key drivers were specific to rosiglitazone, consistent with its broader transcriptional program extending beyond core metabolic reprogramming. Rosiglitazone-unique key drivers included CAV1 and PTPRB in the cytoskeleton and cell mechanics module; TIMP1 and FBN1 in the ECM module; and ADAM8, RTP4, and SOCS3, in the immune signaling module.

**Figure 6.**
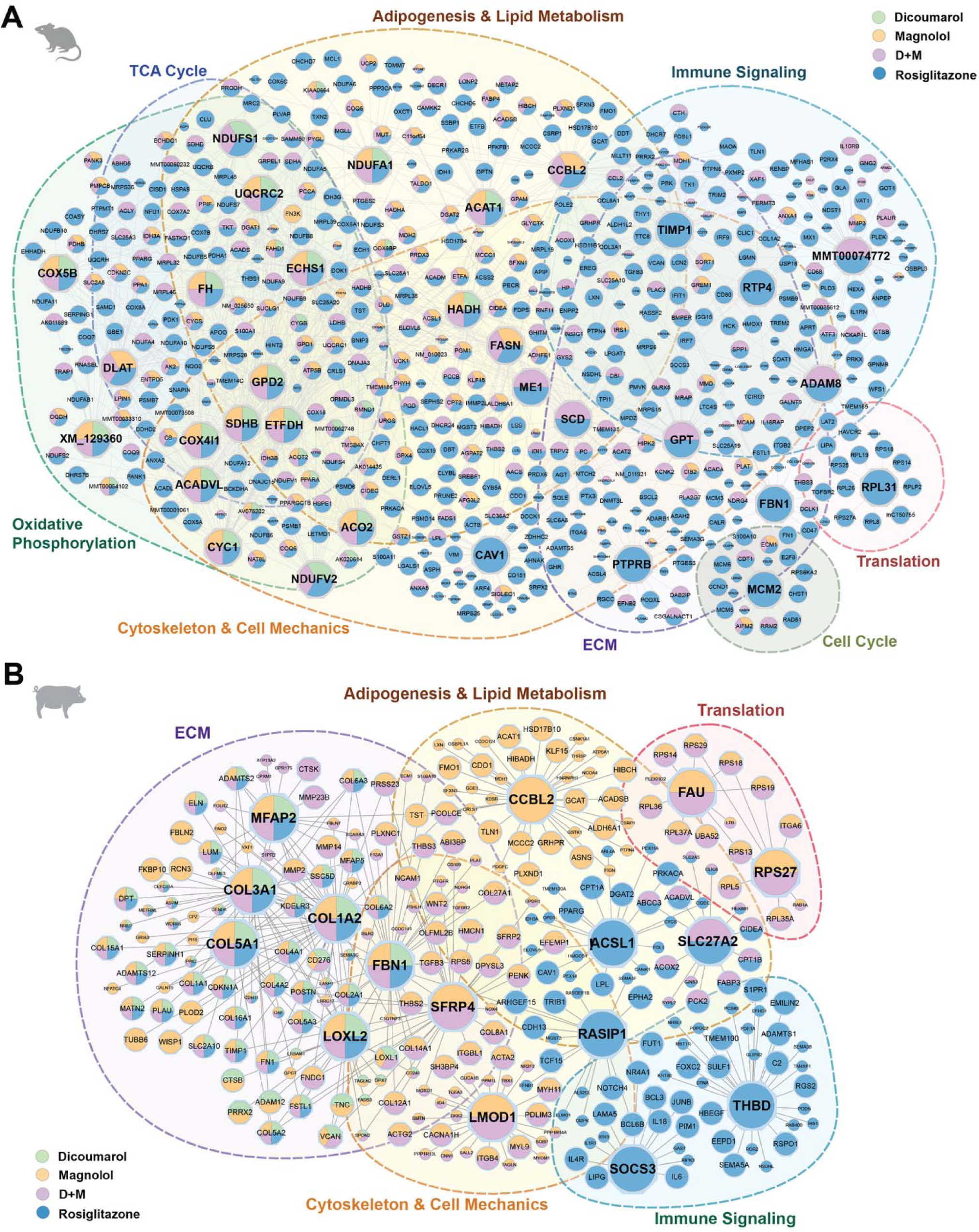
Multi-omics network analysis predicts gene regulatory networks mediating treatment responses across species. (A) Predicted key driver genes and their associated differentially expressed genes (DEGs) across compound treatments constructed from transcriptomic data using Mergeomics. Diamond nodes indicate predicted key drivers (KDs); circular nodes represent other network genes. Node size reflects KD rank. Grey lines present edges denote network connectivity. Pie chart colors indicate treatment contributions (green: dicoumarol, orange: magnolol, purple: D+M, blue: rosiglitazone). For smaller non-KD nodes, colors represent DEGs in each treatment; for KD nodes, node colors indicate the treatment conditions in which gene was identified as a predicted KD in the Mergeomics analysis. Dashed ellipses delineate functional modules: oxidative phosphorylation, TCA cycle, fatty acid β-oxidation, adipogenesis and lipid metabolism, ECM, immune signaling, cytoskeleton and cell mechanics, and cell cycle/translation. (B) Key driver network for porcine DFAT cells, constructed from transcriptomic data using Mergeomics. Node and edge representation as in (A). Dashed ellipses delineate functional modules: ECM, adipogenesis and lipid metabolism, cytoskeleton and cell mechanics, immune signaling, and translation.

Porcine networks exhibited similar functional themes: adipogenesis and lipid metabolism, ECM, cytoskeleton and cell mechanics, translation, and immune signaling (**Figure 6B and Figure S1**). Consistent with cytoskeletal pathway enrichment observed in the DEG analysis (**Figure 5G**), the cytoskeleton and cell mechanics module, which contains the key driver LMOD1, was D+M- and magnolol-weighted, and RASIP1, which was rosiglitazone-weighted. The ECM module was dominated by magnolol and D+M contributions, featuring collagens and ECM proteins such as COL3A1, COL1A2, COL5A1, FBN1, and LOXL2. The adipogenesis and lipid metabolism module showed broader treatment contributions driven by D+M and rosiglitazone, and the drivers included ACSL1, CCBL2, and SLC27A2. The immune signaling module was more specific to rosiglitazone and contained key drivers SOCS3 and THBD.

Comparison of key drivers across murine and porcine cell systems revealed largely shared regulatory programs, including adipogenesis and lipid metabolism; immune signaling; translation; cytoskeleton and cell mechanics; and ECM remodeling (**Figure 6 and Figure S1**). Among the most highly connected predicted key drivers, CCBL2 emerged as a prominent hub in both species; while primarily known for its role in kynurenine metabolism, the consistent emergence of CCBL2 across treatments suggests broader amino acid metabolic remodeling that accompanies adipogenic differentiation^45^. Other shared key drivers included ECM genes such as FBN1, COL1A2, and COL3A1, although these showed different weightings that depended on the treatment: these ECM genes were predominantly rosiglitazone-induced in mouse, but also distributed across magnolol, D+M, and rosiglitazone in porcine. These findings are consistent with transcriptomic data showing ECM remodeling is a rosiglitazone-specific response in 3T3-L1 adipocytes but more broadly implicated across treatments in DFAT cells.

Beyond shared features, we also observed several unique regulatory themes that differed between murine and porcine systems. For example, murine-specific functional themes included TCA cycle, oxidative phosphorylation, and cell cycle programs, reflecting coupling of mitochondrial oxidative metabolism with adipogenesis and lipid uptake in mice. By contrast, porcine cells showed a larger proportion of ECM-related key drivers, demonstrating stronger themes of ECM remodeling and structural reorganization. While individual key drivers within shared themes differed across systems, the overall pattern supports a conserved metabolic core that underlies the compound-induced adipogenesis in both species.

Together, these findings demonstrate that magnolol and dicoumarol enhance adipogenesis across murine, porcine, and ovine systems, with transcriptomic analyses in murine and porcine cells showing that this enhancement involves shared PPARγ-driven transcriptional programs with a more targeted profile than rosiglitazone.

## DISCUSSION

Here, we identify magnolol and dicoumarol as natural adipogenic enhancers that act in concert to promote lipid accumulation across murine, porcine, and ovine adipocytes. Our omics-driven mechanistic studies revealed that, unlike the potent PPARγ agonist rosiglitazone, which is commonly used to drive adipogenesis in cell culture and has broad impact on adipogenesis, metabolic regulation, and immune signaling, magnolol and dicoumarol induce a more focused transcriptional response targeting adipogenesis. Collectively, these findings establish both a screening strategy and a mechanistic framework for developing food-compatible adipogenic media and scalable biomanufacturing of adipose tissue with applications spanning from future food to adipose tissue engineering more broadly.

Dicoumarol and magnolol enabled substantial reductions in synthetic adipogenic inducers while maintaining robust adipogenesis across edible species. The D+M combination supported adipogenesis at levels similar to rosiglitazone treatment. Compared to the IBMX concentration reported to support adipogenesis in porcine DFAT cells (∼500 µM)^6^, D+M reduced IBMX requirements by ∼40-fold (to 12.5 µM). In ovine cells, D+M treatment supported robust adipogenesis with complete elimination of IBMX. These media simplifications directly address major cost and sustainability barriers in cultivated meat production. Specialized media components – including serum, growth factors such as insulin and insulin-like growth factor 1 (IGF-1), and recombinant signaling proteins such as bone morphogenetic protein 4 (BMP4) – require energy- and water-intensive upstream manufacturing and are thus major contributors to both the production costs and environmental footprint of cultivated meat^1,46,47^. While recent cultivated fat protocols have demonstrated adipogenesis in livestock cells using low-serum or serum-free conditions, these still rely on rosiglitazone as a key inducer^6,15,48^. Natural alternatives such as dicoumarol and magnolol that reduce synthetic inducer requirements provide a strategy for more efficient and sustainable engineered adipose tissue production.

Importantly, dicoumarol and magnolol produced substantial adipogenic activity with a more focused transcriptional profile that could provide advantages over synthetic, pharmaceutical-grade thiazolidinediones such as rosiglitazone, which is an insulin-sensitizing drug with a clinically documented side-effect profile^14^. The combined D+M treatment provides similar adipogenic activation to rosiglitazone, and the omics profiles of dicoumarol and magnolol suggest they operate to enhance adipogenesis through complementary pathways. Magnolol exhibited stronger adipogenic transcriptional signatures consistent with partial PPARγ agonism; this mechanism is supported by prior reports of PPARγ activation by magnolia-derived biphenolic compounds, including the isomer honokiol^49,50^. Additional evidence that magnolol alters PPARγ-dependent metabolic processes comes from livestock studies: in laying hens, magnolol supplementation enriches the PPAR signaling pathway and shifts lipid metabolism from fatty acid synthesis toward oxidation^50,51^, indicating that magnolol modulates PPARγ-dependent metabolic programs *in vivo* across species. By contrast, dicoumarol enhanced adipogenesis without dramatic changes in individual genes, yet showed consistent PPAR pathway enrichment across murine and porcine cell systems, consistent with its reported activity as a weak partial PPARγ agonist^52^; these findings suggest that dicoumarol engages adipogenic programs through subtle transcriptional shifts rather than broad transcriptional remodeling. As a vitamin K antagonist that inhibits vitamin K epoxide reductase, dicoumarol may influence adipogenesis through vitamin K-dependent pathways, altered redox and inflammatory signaling^53^, or the coagulation and complement cascades^54^, which are known modulators of adipocyte differentiation^55,56^. In porcine DFAT cells, dicoumarol downregulated redox homeostasis genes including CAT and TXNRD1, suggesting that dicoumarol’s adipogenic effect may be partially mediated through suppression of oxidative stress. Although dicoumarol is a well-characterized NQO1 inhibitor of NAD(P)H quinone dehydrogenase 1 (NQO1) that regulates lipid accumulation during 3T3-L1 adipogenesis^57^, NQO1 transcript levels were not significantly altered in our transcriptomic analysis, suggesting NQO1 inhibition may not be the primary mechanism driving the observed adipogenic effects. Despite reports that the related anticoagulant warfarin inhibits PPARγ signaling in prostate tissue^58^, we show that dicoumarol combined with magnolol achieved pro-adipogenic effects; a potential mechanism could involve complementary targeting of upstream regulators or metabolic nodes that, together with weak or partial PPARγ engagement by both compounds, converge on adipogenic activation. Beyond PPARγ-related mechanisms, both compounds suppressed inflammatory pathways, specifically TNFα/NF-κB signaling, a well-established inhibitor of adipocyte differentiation^59^, suggesting a shared anti-inflammatory axis may additionally contribute to their adipogenic effects.

Network analysis revealed shared and distinct features of regulatory architectures between mouse and porcine systems that have implications for biomanufacturing strategies. Mouse 3T3-L1 adipocytes showed predicted key driver genes that converged on metabolism and lipid-handling hubs across all treatments, consistent with our key finding of dicoumarol and magnolol as adipogenic enhancers with similar effects as rosiglitazone. Porcine DFAT cells also showed activation of core adipogenic pathways but showed stronger modulation of networks with ECM and cytoskeletal regulatory programs, particularly with magnolol treatment. The differences in the proportions of modules that emerge between murine and porcine cells likely reflects the distinct cellular contexts: immortalized 3T3-L1 cells passaged extensively on rigid plastic may have reduced mechanosensing capacity^60^, whereas primary-derived porcine cells retain ECM-responsive differentiation programs. Insights from mechanobiology could have practical relevance for cultivated meat production, since mechanical cues influence differentiation efficiency and tissue structure^61^

More broadly, our findings across murine and porcine cell systems highlight that some core findings – such as regulation of key adipogenic pathways – can translate robustly to edible species, but not all findings from mouse models can predict livestock cell behavior. Notably, the murine cell model showed stronger signs of TCA cycle, oxidative phosphorylation, and fatty acid beta-oxidation, whereas the porcine cells showed more prominent signature of ECM regulation. In porcine cells, magnolol-containing treatments prominently supported collagen and ECM remodeling modules, while rosiglitazone showed inflammatory programs. It is important to note that cross-species comparisons may be constrained by the lack of annotation across species. For example, due to the lack of complete ortholog mapping between porcine and human genomes, we excluded porcine DEGs without human orthologs from pathway enrichment and network analyses; therefore, any biological processes represented by these unmapped genes remain uncharacterized. Improved annotation and orthology mapping will be important for more complete mechanistic understanding across species. While we demonstrated efficacy in porcine and ovine systems, future studies across additional livestock species – including bovine and avian – will be important for establishing broader applicability and mapping the full landscape of shared and unique regulatory pathways relevant to food production.

Both dicoumarol and magnolol are positioned as practical alternatives to pharmaceutical-grade synthetic agonists, which face regulatory barriers and supply chain constraints in food applications. However, translating these findings to food production requires addressing safety, regulatory, and manufacturing challenges. Magnolol has a favorable profile as a food ingredient^28^, and has been explored as a dietary feed supplement in livestock species including pigs, laying hens, and ducklings, where it improves growth performance, gut health, and lipid metabolism without adverse effects^29–32^. The short half-life of magnolol (∼2.33 h in rats)^62^ suggests rapid clearance that could minimize residues in final products. Dicoumarol is a vitamin K antagonist used clinically as an anticoagulant for over 80 years^53,54^, which presents a more complex safety profile due to reported hemorrhagic toxicity at anticoagulant doses^54^. Although both compounds could function as upstream additives to culture medium rather than direct food ingredients, comprehensive safety evaluation will be essential. As with any medium component, regulatory approval pathways will require demonstrating that residual levels pose negligible risk. Beyond safety, practical implementation will require systematic optimization of concentrations and treatment timing across cell types, validation in bioreactor systems at manufacturing scale, and assessment under process conditions that may include variable oxygen, pH fluctuations, and fluid shear stress.

This work establishes a framework for discovering natural compounds that can simplify media formulations while achieving desired cellular outcomes in biomanufacturing applications. By demonstrating that natural bioactive compounds can replace pharmaceutical-grade synthetic agonists while reducing dependence on conventional inducers, we address a key sustainability challenge in cultivated meat: the environmental footprint of upstream reagent manufacturing. Life cycle analyses identify specialized media components as major contributors to the resource intensity of cellular agriculture^47,63^, suggesting that natural compound substitution and media simplification could provide sustainability benefits. The narrower transcriptional profiles of dicoumarol and magnolol compared to synthetic agonists may also provide advantages beyond cultivated meat. In tissue engineering and regenerative medicine contexts where maintaining consistent cellular phenotypes is important, such as generating organoid models or engineering implantable tissues, compounds that activate core differentiation programs with more targeted transcriptional effects may be preferable to inducers that result in more extensive gene expression changes. The screening approach presented here combines high-throughput phenotypic discovery with omics validation in tractable model systems, which provides a strategy for identifying natural compounds that can reduce reliance on synthetic inputs in biomanufacturing applications from cellular agriculture to pharmaceutical production to tissue engineering.

## EXPERIMENTAL PROCEDURES

### Cell culture

Mouse embryonic fibroblast cells (3T3-L1, ATCC, CL-173) were cultured in Dulbecco’s Modified Eagle’s Medium (DMEM) (Gibco, 11995065) supplemented with 10% fetal bovine serum (FBS) (GemCell, 100-500) and 1% antibiotic-antimycotic (Gibco, 15240062) in a 37°C, 5% CO_2_ incubator. The identity of 3T3-L1 cells was confirmed using Short Tandem Repeat (STR) profiling. De-differentiated porcine cells (DFAT) were a generous gift from the Kaplan Lab, Tufts University. Porcine DFAT cells were cultured in DMEM with 20% FBS and 100 µg/ml Primocin (InvivoGen, ant-pm-1). To passage, both murine and porcine cells were trypsinized using 0.25% Trypsin-EDTA (Thermo Fisher, 25200-056). Primary ovine preadipocytes (Opo Bio, Opo-Baa-A17) were cultured in Dulbecco’s Modified Eagle Medium/Nutrient Mixture F-12 (DMEM/F12) (Gibco, 11320033) supplemented with 10% FBS and 1% antibiotic-antimycotic, and passaged using TrypLE Express Enzyme (Gibco, 12604013). 3T3-L1 and ovine cells were cultured for no more than 10 passages, and porcine DFAT cells were cultured for no more than 15 passages. Cells were cryopreserved in culture medium with 10% v/v DMSO (Supelco, MX1456P-6). Cell cultures were routinely screened for mycoplasma contamination using the Mycoplasma PCR Detection Kit (Applied Biological Materials, G238).

### High-throughput screen to identify adipogenic enhancers

To identify adipogenic enhancers, we seeded 3T3-L1 cells in 384-well microplates (Greiner, 781096) at a density of 3,400 cells per well (>90% confluence) using a multiwell dispenser (MultiDrop, Thermo Scientific). To disentangle effects of the top compounds on promoting adipogenesis versus cell proliferation, we seeded cells at >90% confluency in differentiation media. We established the screen to identify compounds that increase lipid accumulation compared to the vehicle control, dimethyl sulfoxide (DMSO) (Supelco, MX1456P-6), by culturing cells in a commercial adipocyte differentiation medium (iXCells, MD-0005) that was diluted to 7.5% v/v in DMEM. Compounds from the Spectrum Collection (MicroSource), Natural Organic Compound Library (Selleckchem, L7600), and Natural Product Library (Selleckchem, L1400) were dissolved in the solvent recommended by the manufacturer, either DMSO or water. As a positive control, we treated cells with the PPARγ agonist, rosiglitazone (1 µM; TCI, R0106). For experimental consistency, the final concentration of DMSO in the culture medium was adjusted to 0.1% (v/v) for all treatment and control groups. After incubating the treated cells for 4 days at 37 °C in a humidified incubator with 5% CO_2_, we measured lipid accumulation by visualizing intracellular lipids with BODIPY 493/503 (Cayman Chemical, 25892) and DNA with Hoechst 33342 (Invitrogen, H3570), and conducting quantitative image analysis (as described below).

### Validating top hits

For validation experiments, we created stock solutions of dicoumarol (10 mM, Cayman Chemical, 20764) and magnolol (100 mM, Cayman Chemical, 528438) in DMSO. Rosiglitazone was used at 1 µM as a positive control. The final concentration of DMSO in the culture medium was maintained at 0.1% (v/v) for individual drug treatments and 0.2% (v/v) for co-treatment experiments, with corresponding vehicle controls adjusted accordingly. To validate the effects of magnolol and dicoumarol on adipogenesis of 3T3-L1 cells, we induced differentiation using DMEM containing magnolol or dicoumarol and 0.75 µg/mL insulin (Sigma-Aldrich, I0516), 75 nM dexamethasone (Sigma-Aldrich, D4902), and 37.5 µM isobutyl methylxanthine (IBMX, Sigma-Aldrich, I5879) with 10% FBS and 1% antibiotic-antimycotic. We also tested the effects of dicoumarol and magnolol on DFAT cells differentiated in Advanced DMEM (Gibco, 12491015) supplemented with 100 µg/ml Primocin, 0.5 µM dexamethasone, 12.5 µM IBMX, 2 mM (1×) GlutaMAX (Thermo Fisher, 35050-061), 20 µM biotin (TCI America, B04631G), 10 µM calcium-D-pantothenate (TCI America, P001225G), and 2% FBS. We additionally tested dicoumarol and magnolol on ovine preadipocytes in Advanced DMEM media containing 0.5 µM dexamethasone, 2 mM (1×) GlutaMAX, 20 µM biotin, and 10 µM calcium-D-pantothenate with 2% FBS and 1% antibiotic-antimycotic. Media was changed every two days during differentiation.

### Viability assay

Cell viability was assessed using a live/dead assay with Calcein-AM (Invitrogen, C1430) and propidium iodine (PI; Invitrogen, P3566). At the indicated treatment timepoints, a staining solution was added to each well of the culture plates to achieve final concentrations of Calcein-AM (5 µM) and PI (1 µg/mL). Cells were incubated for 15 minutes at 37°C protected from light, then imaged and viability quantified as described below.

### Microplate fluorescence imaging and quantification

Microplate fluorescence imaging was performed using an automated confocal microscope outfitted with a 20× objective (N.A. 0.75) (Molecular Devices); images were analyzed using MetaXpress software (Molecular Devices). For lipid accumulation assays, intracellular lipids were visualized with BODIPY 493/503 (FITC filter set) and nuclei were counterstained with Hoechst 33342 (DAPI filter set); lipid area per cell was quantified by normalizing total lipid area to the number of nuclei using the Granularity module in MetaXpress. For viability assays, live cells were stained using Calcein-AM (FITC filter set), and dead cells were stained with PI (TexasRed filter set). Cell viability was calculated as the percentage of Calcein-positive cells relative to total Calcein- and PI-positive cells using the Live/Dead application module in MetaXpress.

### Suspension culture of adipogenic microtissues

To test the efficacy of dicoumarol and magnolol on promoting adipogenesis in tissues cultured using a scalable approach that is compatible with food applications, we cultured adipocytes on edible microcarriers in suspension culture. In this culture system, cells and microcarriers form microtissue aggregates over the experimental timescale of 7 days. We fabricated edible gelatin microcarriers using water-in-oil emulsions and cultured the tissues in a multiwell plate on a nutator as previously described^34,35^. To minimize cell adhesion, we pre-treated multiwell plates prior to seeding cells with sterile 0.2% Pluronic solution for 10 minutes, followed by aspiration, air drying, and three washes with 1×PBS. We seeded preadipocytes on the microcarriers at a density of 26,400 cells per 0.88 cm² of microcarriers in 100 µL of culture media in each well. To induce differentiation, we replaced the medium with differentiation induction media after 48 hours. Both ovine and porcine cells were differentiated for 7 days, after which the resultant adipogenic microtissues were collected, fixed with 4% paraformaldehyde (Thermo Scientific, J19943K2), and stained with BODIPY and Hoechst 33342 for confocal microscopy.

### Confocal imaging and analysis of adipogenic microtissues

To visualize lipid accumulation in adipogenic microtissues, we imaged the fixed and stained microtissues using a confocal laser scanning microscope (Zeiss LSM 880) with a 10× objective (N.A. 0.45). Z-stack images were acquired with a step size of 25 µm across five optical sections (optical section thickness ∼3.8 µm for DAPI channel and ∼4.3 µm for FITC channel, estimated from N.A. and emission wavelength), with the pinhole set to 1 Airy unit. For quantitative image analysis, we used the Bio-Formats toolbox in MATLAB. Nuclei were segmented in each Z-slice following intensity adjustment, background subtraction, Gaussian smoothing (σ = 3), adaptive thresholding (sensitivity = 0.4), and removal of small objects (<90 pixels). To quantify lipid accumulation, we measured the total fluorescence intensity per slice. We normalized fluorescence intensity values per nucleus to extract a lipid intensity per cell and lipid area per cell.

### RNA-sequencing (RNAseq)

To determine transcriptomic changes induced by dicoumarol and magnolol, we conducted bulk RNAseq of 3T3-L1 and DFAT adipocytes after 5 days of differentiation (n = 3/group). Total RNA was extracted and quality-checked using the Agilent TapeStation, and processed by the UCLA Technology Center for Genomics & Bioinformatics; only samples with RNA Integrity Number (RIN) ≥ 7.0 were used for library preparation. Paired-end sequencing of the cDNA (2×150 bp, 30 million reads per sample) was conducted on the Illumina NovaSeq X Plus platform. Reads were aligned using STAR with mouse reference genome GRCm38 (Ensembl release 107) and pig reference genome sus scrofa 11.1 (Ensemble release 110). Gene counts were normalized and differential gene expression was assessed using DESeq2. P-values were adjusted using the Benjamini-Hochberg false discovery rate (FDR), and genes with FDR < 0.05 were considered differentially expressed.

Gene set enrichment analysis (GSEA) was performed using the clusterProfiler (v4.18.4) and enrichplot (v1.30.5) packages in R, with genes pre-ranked by a signed ranking metric calculated as -log_10_(p-value)×sign(log_2_(fold change)). For porcine data, gene identifiers were mapped to human ortholog gene symbols prior to enrichment analysis to enable use of human-curated gene set collections. Enrichment was assessed against the Hallmark pathways, Gene Ontology Biological Process (GO BP), and Kyoto Encyclopedia of Genes and Genomes (KEGG) gene set collections from the Molecular Signatures Database (MSigDB v2024.1), accessed via msigdbr (v26.1.0). Mouse and porcine GO:BP annotations were sourced from org.Mm.eg.db (v3.22.0) and org.Ss.eg.db (v3.22.0) respectively; KEGG pathway annotations were accessed via the KEGG REST API in April 2026. Gene sets with p_adj_ < 0.05 were considered significantly enriched. Overrepresentation analysis (ORA) of DEGs was performed using the clusterProfiler against the same Hallmark, GO BP, and KEGG gene set collections.

### Mergeomics analysis

To identify enriched pathways, networks and potential network regulators (termed key drivers) in transcriptional changes, we applied Mergeomics^44^. Mouse and porcine gene identifiers were automatically mapped to human ortholog gene symbols within the Mergeomics pipeline prior to network analysis. P-values derived from differential expression analysis (RNA-seq) were used as input for Marker Set Enrichment Analysis (MSEA), which identified pathways and gene sets that were significantly enriched. The top 15 enriched modules from MSEA were subsequently mapped onto a Bayesian adipose tissue network in Mergeomics, which was constructed from tens of human and mouse multiomics datasets. Weighted key driver analysis (wKDA) was performed on this network to identify central regulatory genes within the enriched modules, highlighting potential upstream drivers of the observed molecular signatures. We used Cytoscape (version 3.10.4) to visualize the compiled networks of all treatment groups.

## RESOURCE AVAILABILITY

### Lead contact

Further information and requests for resources and reagents should be directed to and will be fulfilled by the lead contact, Amy Rowat.

### Materials availability

This study did not generate new unique reagents.

### Data and code availability

All data reported in this paper will be shared by the lead contact upon request. The RNAseq data generated in this study are available in the NCBI Expression Omnibus (GEO) under accession number GSE336373. This paper does not report original code.

## ACKNOWLEDGMENTS

This work was conducted through support from a Noble Family Innovation Fund Seed Grant (to ACR and RD), the State of California, and the National Science Foundation (BRITE Fellow Award to ACR). We are grateful to Amanda Faye Lipsey, of Amanda Faye Consulting, for her support in preparing this manuscript and the Technology Center for Genomics & Bioinformatics (TCGB) at UCLA for providing sequencing, library preparation, and bioinformatics services and support.

## AUTHOR CONTRIBUTIONS

Conceptualization, A.C.R. and R.D.; Methodology, Q.X., N.S.K., K.K.C., C.A.C., R.D., and A.C.R.; Investigation, Q.X., K.K.C., and E.C.; Formal Analysis, Q.X.; Data Curation, Q.X. and E.C.; Software, M.B. and X.Y.; Visualization, Q.X., R.D., and A.C.R.; Writing – Original Draft, Q.X. and A.C.R.; Writing – Review & Editing, Q.X., N.S.K., C.A.C., E.C., M.B., X.Y., R.D., and A.C.R.; Supervision, X.Y., R.D., and A.C.R.; Funding Acquisition, A.C.R.; Resources, M.B., X.Y., R.D., and A.C.R.; Project Administration, A.C.R.

## SUPPLEMENTAL MATERIALS

## SUPPLEMENTAL MATERIALS

**Figure S1.**
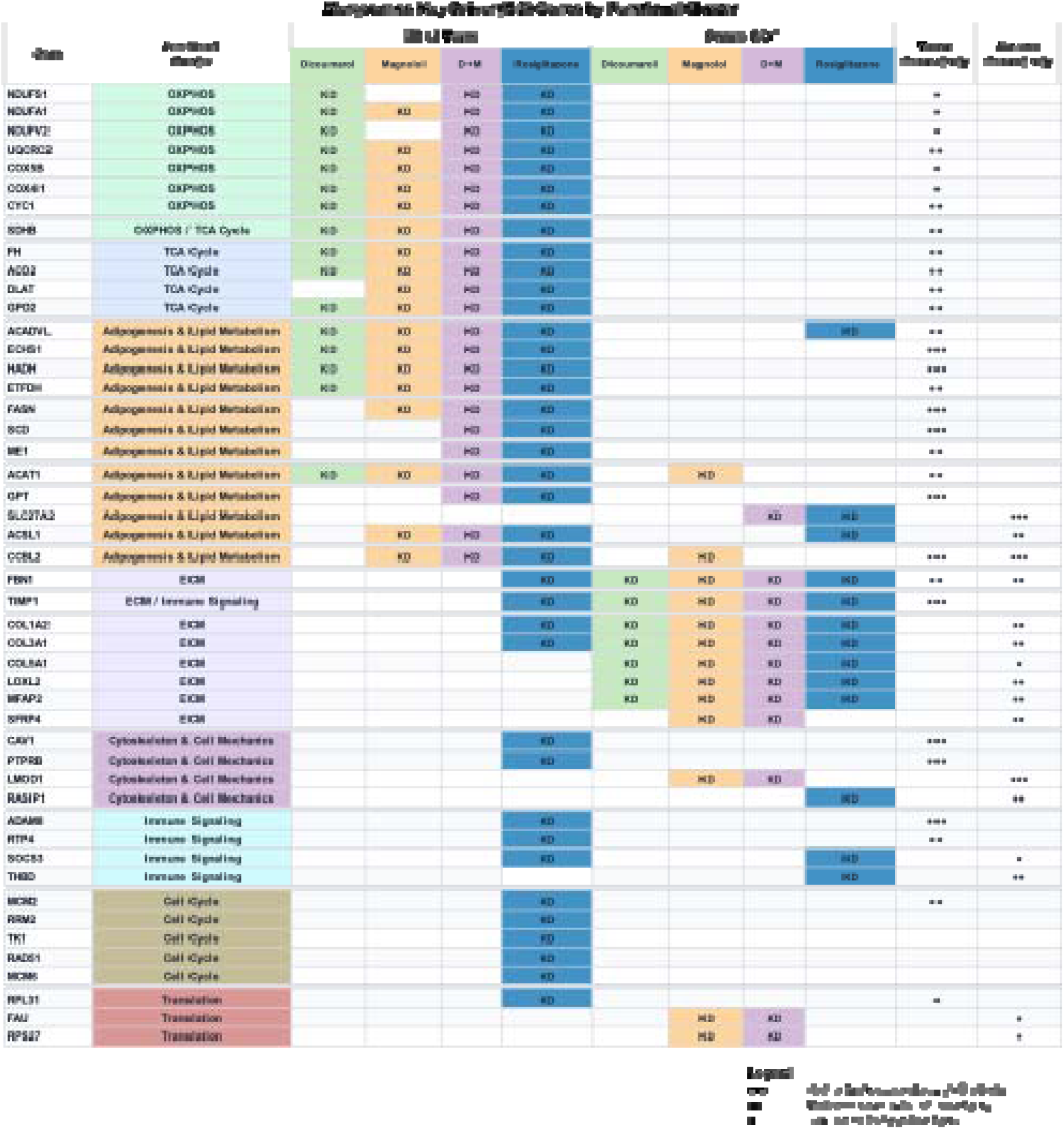
Predicted key driver (KD) genes from Mergeomics network analysis summarized by functional cluster. Gene identified as key drivers in the Mergeomics analysis of 3T3-L1 adipocytes and DFAT cells are listed with their functional cluster assignments and treatment-specific key driver status. Colored cells indicate that the genes were identified as a predicted key driver in that treatment condition for the indicated cell systems (Dicoumarol, Magnolol, D+M, or Rosiglitazone). Network connectivity reflects the number of neighbor genes connected to each key driver node in the Mergeomics network: ••• high (>60 edges); •• medium (30-60 edges), • low (<30 edges). Functional clusters were assigned based on gene function and Mergeomics module annotations and correspond to the cluster annotations shown in Fig. 6. Genes belonging to multiple functional themes are annotated with combined cluster labels.

